# Integrin αvβ8 on T cells is responsible for suppression of anti-tumor immunity in multiple syngeneic models and is a promising target for tumor immunotherapy

**DOI:** 10.1101/2020.05.14.084913

**Authors:** Eswari Dodagatta-Marri, Hsiao-Yen Ma, Benjia Liang, John Li, Dominique S. Meyer, Kai-Hui Sun, Xin Ren, Bahar Zivak, Michael D. Rosenblum, Mark B. Headley, Lauren Pinzas, Nilgun I. Reed, Joselyn S. Del Cid, Maeva Adoumie, Byron C. Hann, Sharon Yang, Anand Giddabasappa, Kavon Noorbehesht, Bing Yang, Joseph Dal Porto, Tatsuya Tsukui, Kyle Niessen, Amha Atakilit, Rosemary J. Akhurst, Dean Sheppard

**Affiliations:** Helen Diller Family Comprehensive Cancer Center, University of California, San Francisco, CA, USA; Lung Biology Center, Department of Medicine, University of California, San Francisco, CA, USA; Department of Gastrointestinal Surgery, Shandong Provincial Hospital Affiliated to Shandong University, Jinan 250021, Shandong, China; Department of Dermatology, University of California, San Francisco, CA, USA; Department of Pathology, University of California, San Francisco, CA, USA; Comparative Medicine, Pfizer Inc., San Diego, CA, USA; Oncology Research Unit, Pfizer Inc., Pearl River, NY, USA; Pfizer Centers for Therapeutic Innovation, San Francisco, CA, USA; Department of Anatomy, University of California, San Francisco, CA, USA

## Abstract

The αvβ8 integrin is a key activator of transforming growth factor β (TGF β), which has been shown to inhibit anti-tumor immunity. Previous work has suggested that αvβ8 on tumor cells could modulate tumor growth and responses to immune checkpoint blockade. We now show that a potent blocking monoclonal antibody against αvβ8 (ADWA-11) causes growth suppression or complete regression in syngeneic models of squamous cell carcinoma (CCK168), mammary cancer (EMT-6), colon cancer (CT26), and prostate cancer (TRAMPC2), especially when it is combined with other immunomodulators (anti-PD-1, anti-CTLA-4 or 4-1BB) or radiotherapy. αvβ8 is expressed on tumor cells in some of these models, but tumor cell expression of αvβ8 is not essential for the beneficial effects of ADWA-11 therapy. αvβ8 is consistently expressed at highest levels on CD4+CD25+ T cells within tumors, and specific deletion of *Itgb8* from T cells is as effective as ADWA-11 in suppressing tumor growth. Treatment with ADWA-11 increases expression of a suite of genes in tumor infiltrating CD8+ T cells that are normally inhibited by TGFβ and are involved in tumor cell killing, including Granzyme B and Interferon-γ. These findings solidify αvβ8 integrin as a promising target for cancer immunotherapy, even for tumors that do not express this integrin.

## Introduction

Immune checkpoint inhibitors have revolutionized the treatment of cancer by allowing host adaptive immunity to eliminate tumor cells. For example, antibodies targeting the immune checkpoint, PD-1, or its ligand, PD-L1 can induce persistent anti-tumor immunity and have become standard therapies for melanoma, lung cancer, head and neck cancers, renal cell carcinoma and bladder cancer(Sharma and Allison, 2015). However, only a minority of affected patients benefit from these treatments. Therefore, intense efforts are underway to develop additional immunomodulatory strategies to extend the reach of this exciting new approach to treating cancer.

Transforming growth factor β (TGFβ) is a potent suppressor of adaptive immunity and an important mediator of immune suppression by a subset of regulatory T cells. TGFβ also promotes secretion and accumulation of a fibrotic tumor stroma that has been proposed as a possible contributor to exclusion of immune cells from some solid tumors. For all of these reasons, inhibition of TGFβ has been widely explored as an adjunctive immunotherapy(Gorelik and Flavell, 2001).

Previous studies have shown that inhibition of TGFβ signaling can enhance responses to checkpoint inhibitors(Dodagatta-Marri et al., 2019; Mariathasan et al., 2018; Tauriello et al., 2018). However, TGFβ plays important homeostatic roles in many biological systems, so effective systemic targeting of TGFβ signaling would likely present challenges due to unwanted side effects(Akhurst and Hata, 2012; Flavell et al., 2010; Yang et al., 2020). Strategies that limit inhibition of TGFβ to specific biological contexts, especially those that contribute to suppression of tumor immunity, could have significant safety and therapeutic advantages over systemic TGFβ inhibition. One such strategy takes advantage of the fact that TGFβ is synthesized as a latent form that can be activated by binding to specific integrin receptors that are restricted in where and when they perform this function, and that can only activate TGFβ1 and 3 (Robertson et al., 2015). Blockade of integrins, that do not affect activation of TGFβ2 homodimers, may be advantageous in avoiding possible outgrowth of dormant metastatic tumor cells that are growth inhibited by TGFβ2 in bone and lymph nodes(Bragado et al., 2013; Jiang et al., 2019; Yumoto et al., 2016).

One previous study showed that an antibody against αvβ8 could inhibit the growth of syngeneic tumors in mice, but that study suggested that this effect required αvβ8 on tumor cells(Takasaka et al., 2018). In the current study, we utilized a potent αvβ8 blocking monoclonal antibody (ADWA-11) we had previously generated by immunizing *Itgb8* knockout mice with recombinant αvβ8(Stockis et al., 2017), to examine whether inhibition of this integrin could facilitate anti-tumor immunity. This antibody was highly potent in inhibiting αvβ8-mediated TGFβ activation in a co-culture bioassay system, and we found that it potently inhibited the growth of tumors whether or not the tumor cells expressed αvβ8. All responding tumor types showed highest levels of αvβ8 expression on CD25+ CD4+ T cells. Deletion of *Itgb8* specifically in T cells was as effective in suppressing tumor growth as was ADWA-11, and ADWA-11 treatment did not further inhibit tumor growth in mice lacking *Itgb8* in T cells. These results are consistent with the idea that inhibition of αvβ8 enhances anti-tumor immunity by blocking αvβ8-mediated TGFβ activation by T cells, and suggest that inhibition of this integrin could be a promising therapeutic strategy for a wide array of tumors, irrespective of tumor cell expression of αvβ8.

## Results

### Effects of combinatorial therapy of ADWA-11 and anti-PD-1 in CCK168 squamous cell, EMT6 mammary and TRAMPC2 prostate carcinoma models

We began by examining the effects of ADWA-11, alone, or in combination with anti-PD-1, in established syngeneic tumor models of squamous cell carcinoma (CCK168 cells) and mammary carcinoma (EMT6 cells) (Figure 1A), two models in which systemic blockade of TGFβ has been shown to enhance responses to immunotherapy(Dodagatta-Marri et al., 2019; Mariathasan et al., 2018). We injected CCK168 cells subcutaneously into the flanks of syngeneic mice or EMT-6 cells directly into the fourth mammary fat pad, and allowed tumors to grow to 65mm^3^ before beginning antibody therapy. Mice were then injected with ADWA-11 or isotype-matched control antibody on days 0 and 7 and anti-PD-1 or its isotype-matched control antibody on days 0, 4 and 8. Tumor diameter was measured every other day and mice were euthanized when tumors reached a size greater than 2000 mm^3^. CCK168 tumors had minimal responses to anti-PD-1, but most mice treated with ADWA-11 monotherapy showed tumor regression, with 5 of 10 mice showing complete tumor regression (Figure 1B, C). Combination therapy with ADWA-11 and anti-PD-1 induced complete regression in 9 of 10 tumors, with a significant increase in long-term survival. We then examined the effects of ADWA-11 on the EMT6 model of mammary carcinoma, that has an immune excluded tumor microenvironment(Mariathasan et al., 2018) and minimal levels of αvβ8 expression (Figure S1D). Anti-PD-1 alone had no effect in this model, ADWA-11 monotherapy had a modest effect and combination therapy substantially and significantly inhibited tumor growth and increased survival (Figure 1D, E). 8 of 16 mice demonstrated complete regression of the EMT6 tumor after combined treatment with ADWA-11 and anti-PD-1.

**Figure 1.**
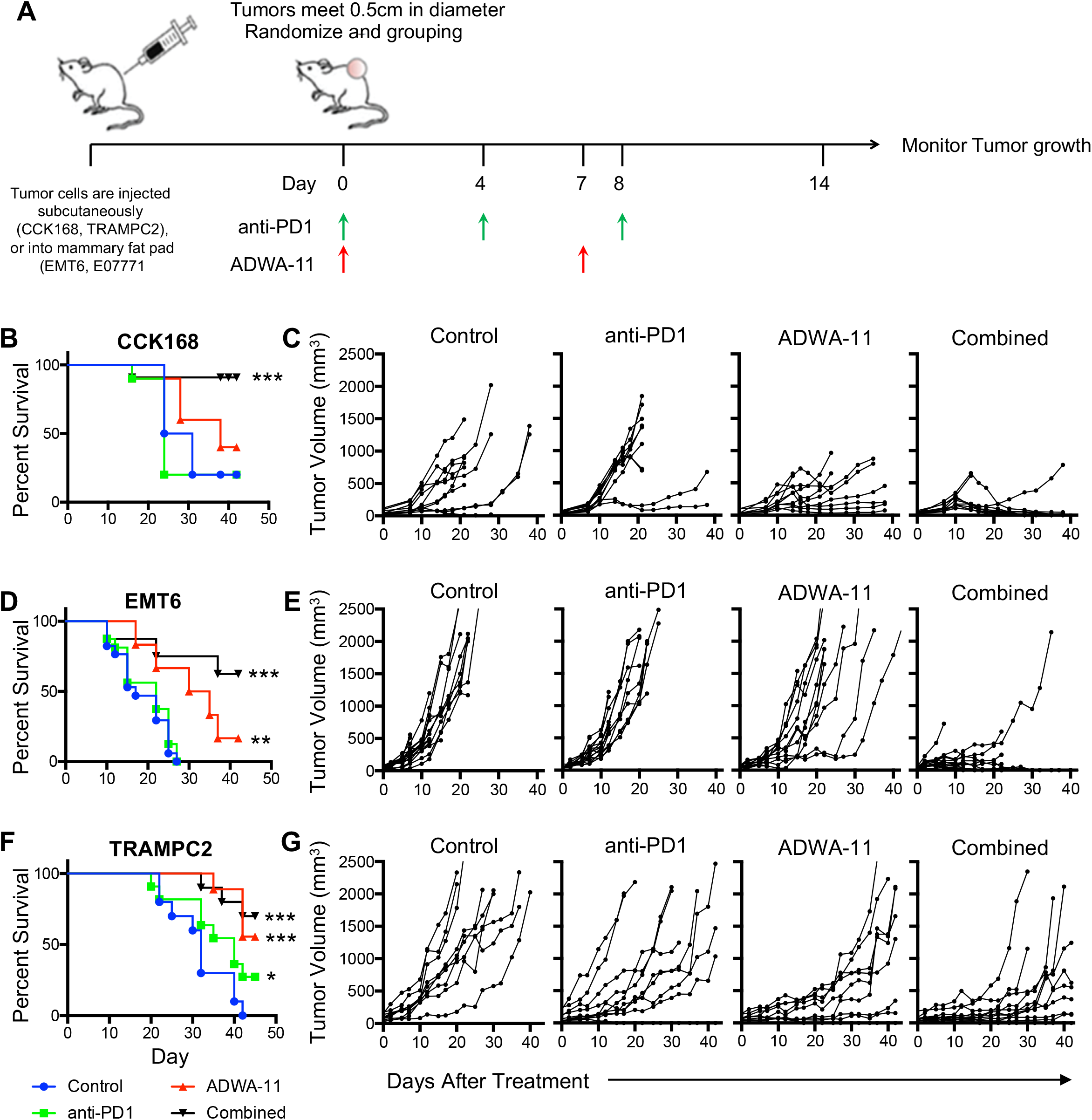
ADWA-11 synergizes with anti-PD-1 to improve survival and decrease tumor size in two solid tumor models. (A) Schematic of multiple tumor models with timeline for injection of tumor cells and antibody injection of 4 treatment groups. Anti-PD-1 was injected i.p. on days 0, 4, and 8 after tumor reached appropriate size for inclusion into the study. ADWA-11 was injected i.p. on days 0 and 7. Kaplan-Meier survival curves (B, D, and F, and individual growth curves of tumors measured every other day for CCK168 (B, C), EMT6 (D, E), and TRAMPC2 (F, G). Mice were euthanized when tumors reached ≥ 2000 mm^3^, if extensive tumor ulceration was observed or at the 45-day endpoint. n=10-18 in each group. *p<0.05, **p<0.01 ***p<0.001 by log-rank Mantel-Cox test.

To take advantage of genetic tools to evaluate functional relevance of αvβ8 - expression in specific cell types, we also examined the effects of ADWA-11, with or without anti-PD-1, in a C57BL/6 syngeneic model, TRAMPC2 prostate cancer. TRAMPC2 is a prostatic adenocarcinoma developed from transgenic mice expressing the SV40 Large T antigen specifically in prostatic epithelium(Foster et al., 1997). Mice harboring subcutaneous syngeneic TRAMPC2 tumors showed minimal response to anti-PD-1 monotherapy, but tumor growth was markedly reduced by treatment with ADWA-11, with no additional effect of inhibition of PD-1 (Figure 1F, G).

### ADWA-11 increases expression of cytotoxic granzyme B in CD8+ T cells and of interferon-γ in both CD4+ and CD8+ T cells

We next characterized the effects of ADWA-11 (with or without anti-PD-1) on the nature of the immune infiltrate(Thomas and Massague, 2005; Wu et al., 2014) in the CCK168 tumor model. Although treatment with anti-PD-1 in this model had no effect on the total number of intratumoral CD8+ T cells, treatment with ADWA-11 significantly increased CD8+ T cell accumulation (Figure 2A). ADWA-11 also significantly increased the percentage of CD8+ T cells that expressed granzyme B, and elevated expression of IFNγ in both CD4+ and CD8+ T cells (Figure 2A), whereas treatment with anti-PD-1 alone had no effect on these endpoints. We were unable to identify significant production of IL-17 in any T cells subset (data not shown). We also performed immunophenotyping of tumors from mice treated with IgG control and ADWA-11 of the EMT6 (Figure 2B) and TRAMPC2 (Figure 2C) models. In each of these tumors, ADWA-11 increased CD8+ T cell accumulation, Granzyme B expression in CD8+ T cells, and IFNγ expression in tumor CD4+ and CD8+ cells. We did not see consistent effects of ADWA-11 on the numbers of CD4+ T cells, macrophages or dendritic cells. These results suggest that ADWA-11 inhibits in vivo tumor growth and enhances the effects of anti-PD-1 primarily by promoting CD8+ T cell accumulation, Granzyme B expression in CD8+ T cells and IFNγ expression in both CD4+ and CD8+ T cells.

**Figure 2.**
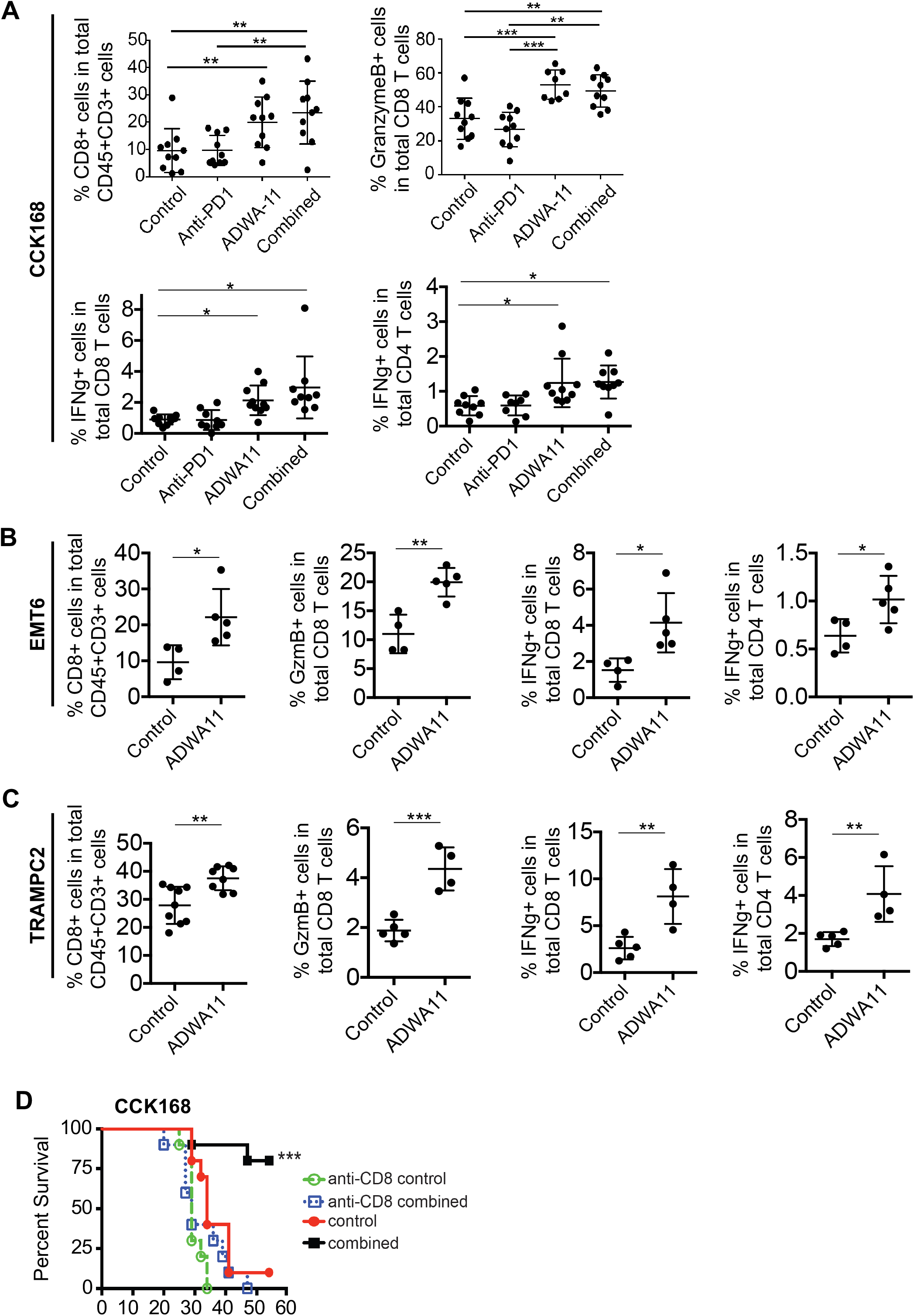
Treatment with ADWA-11 increases accumulation of Granzyme B expressing CD8+ T cells and depletion of CD8+ T cells abolishes its protective effect. (A)Percentage of total CD8+ T cells, granzyme B expressing CD8+ T cells and of IFNγ expressing cells in CD8+ and CD4+ T cells in each mouse with CCK168 tumors treated with control antibodies alone, anti-PD-1, ADWA-11 or combined anti-PD-1 andADWA-11. Flow cytometry was performed to gate these populations (Figure S2). Data in graphs are mean ± S.D, n=10 per group. *p<0.05, **p<0.01, ***p<0.001 by one-way ANOVA. (C) Percentage of IFNγ expressing cells in CD8+ and CD4+ T cells in each mouse in each group. Flow cytometry gating strategy is shown in Figure S2. Data in graphs are mean ± S.D, n=10 per group. *p<0.05 by one-way ANOVA. Representative flow cytometry plots of EMT6 (B) and TRAMPC2 (C) tumors treated with control and ADWA-11, showing CD8+ cells population in total CD45+CD3+ cells, Granzyme B expressing in CD8+ T population, IFNγ expressing in CD4+ and CD8+ T cells. Data in graphs are mean ± S.D, n=5 per group. *p<0.05, **p<0.01, ***p<0.001 by t-test (D) Survival of mice harboring CCK168 tumors pretreated with anti-CD8+ depleting antibody or control antibody one-day prior to initiation of ADWA-11/anti-PD-1 combination therapy. Data reported as percent survival, n=10 in each group. ***p<0.001 by log-rank Mantel-Cox test.

### The beneficial effects of combined ADWA-11 and anti-PD-1 were abrogated by CD8+ T cell depletion

Because the most dramatic effects of ADWA-11 we observed were on CD8+ T cells, we sought to determine whether effects on these cells were responsible for the protective effects of ADWA-11. For this purpose, we treated CCK168 tumor-bearing mice with drugs following removal of CD8+ T cells using a CD8-depleting antibody. Immunostaining showed effective CD8+ T cell depletion (Figure S3), which completely abrogated the beneficial effects of combination therapy with ADWA-11 and anti-PD-1 (Figure 2D and S3).

### Effectorless ADWA-11 inhibits tumor growth, improves survival and induces persistent anti-tumor immunity in CT26 colon cancer and CCK168 SCC

Because our initial studies were undertaken with native murine ADWA-11, we next determined whether any of the protective effects of ADWA-11 could be due to Fc-dependent effects such as antibody-dependent cellular cytotoxicity (ADCC) rather than activation of the host immune response by functionally blocking binding of integrin β8 its ligands. Toward this end, we generated a recombinant, effectorless version of ADWA-11 (ADWA-11_4mut) with 4 mutations in the IgG1 Fc domain that have previously been shown to completely abrogate binding to Fc receptors(Alegre et al., 1992; Young et al., 2014). This effectorless antibody, like wild-type ADWA-11, synergized with anti-PD-1 to enhance tumor survival in the CCK168 model (Figure S4). We used this antibody, in combination with radiotherapy in the CT26 colon carcinoma model. CT26 was tested because it completely lacks detectable αvβ8 expression (Figure S1D) and because radiation therapy has been demonstrated to promote a robust tumor growth inhibition when combined with TGFβ receptor inhibitors (Young et al., 2014). The addition of either ADWA-11_4mut or anti-PD-1 significantly increased survival of mice when combined with radiotherapy (RT) in this model, with ADWA-11 + RT leading to complete regression in 5 of 9 mice (Figure 3A, B). Interestingly, the addition of anti-PD-1 to ADWA-11 added little additional benefit in this model, providing further evidence that inhibition of αvβ8 can be effective even in the absence of additional checkpoint inhibitors. The surviving mice that received either monotherapy or combination therapy were re-challenged with the same tumor cells at >50 days after initial therapy in both the CCK168 and CT26 models. There was minimal tumor growth observed, less than 100 mm^3^, and all tumors that did grow out subsequently regressed despite no further treatment, suggesting that successful treatment with ADWA-11 can lead to long-term anti-tumor immunity, as has been previously described for other immunomodulators, including TGFβ blockade (Figure 3C) (Dodagatta-Marri et al., 2019).

**Fig. 3.**
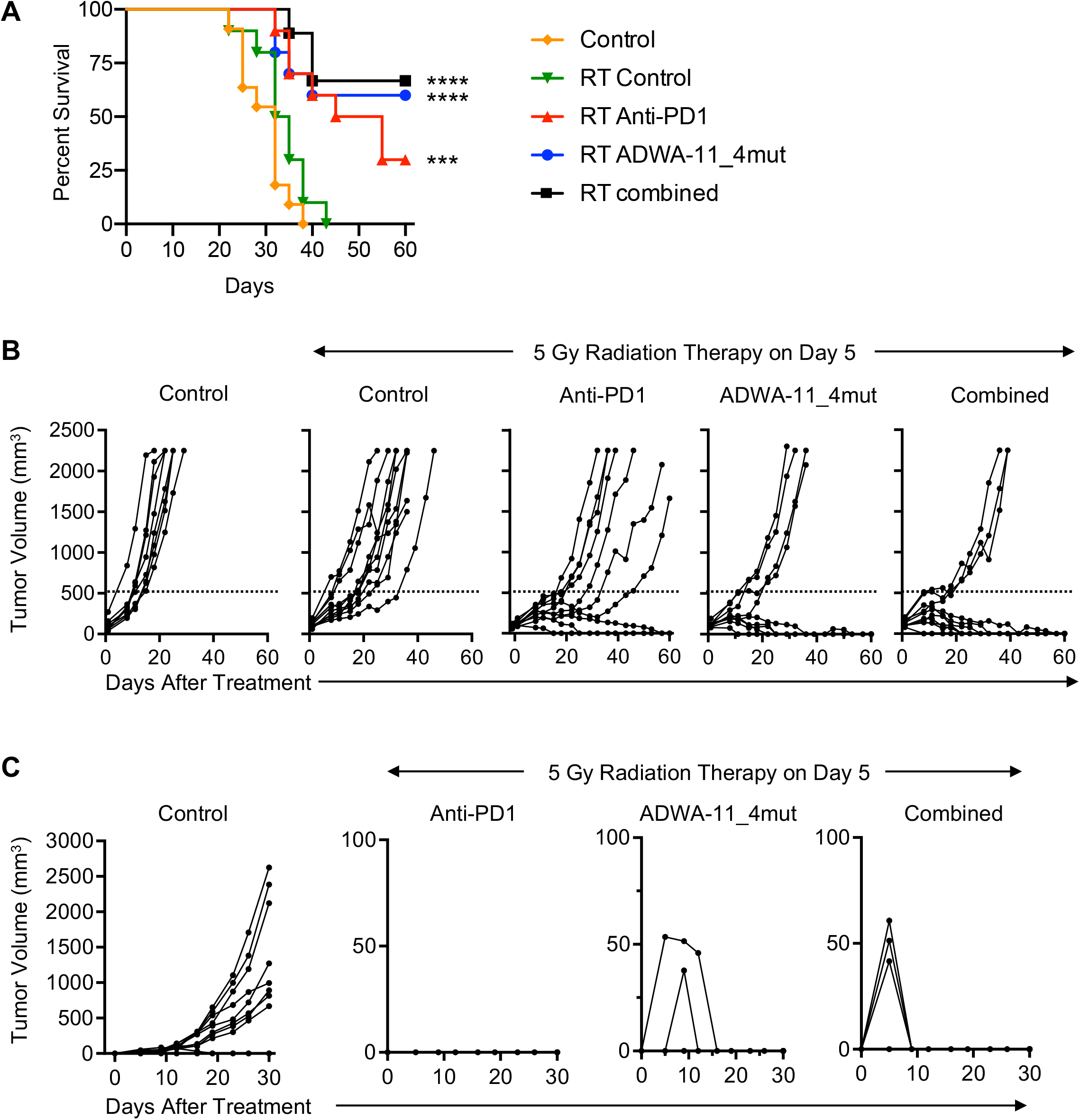
Effectorless ADWA-11_4mut enhances the anti-tumor effects of radiotherapy (RT) and induces long-term anti-tumor immunity in the CT26 colon carcinoma model. (A) Survival curves and (B) individual tumor growth curves in mice implanted with subcutaneous CT26 cells treated with isotype control antibodies, anti-PD-1, ADWA-11_4mut, or a combination of anti-PD-1 and ADWA-11_4mut, plus 5 Gy radiation dose on day 5. One group of mice treated with isotype control antibody did not receive radiation therapy. Data reported as percent survival, n=10 in each group. *** p<0.001, ****p<0.0001 by log-rank Mantel-Cox test. (C) Tumor re-challenge in CT26-cured mice that survived 50 days post-treatment initiation. Parental CT26 cells were implanted into the flank contralateral to that of the original tumor implantation site of CT26-cured mice, 51 days after initiating immunotherapy in combination with radiation therapy. Control mice did not receive radiation and were not previously exposed to tumor cells. Re-challenged mice were followed for 30 days. Control, n= 10; RT plus anti-PD-1, n=3; RT plus ADWA-11_mut, n=5; RT plus ADWA-11_mut and anti-PD-1, n=7.

### ADWA-11_4mut enhances the effects of anti-CTLA-4 and 4-1BB, in the EMT6 mammary carcinoma model

We next sought to determine whether effectorless ADWA-11 might broadly enhance the beneficial effects of additional immunomodulatory therapies. For this purpose, we examined anti-CTLA-4, which has recently be shown to work in part by a different molecular mechanism than anti-PD-1(Wei et al., 2017), and an agonist antibody against 4-1BB, an activator of the inducible costimulatory receptor CD137. We used the EMT-6 model, in which ADWA-11 monotherapy was only minimally effective. In this model, both anti-CTLA4 and anti-4-1BB caused modest reductions in the time to reach the terminal endpoint of tumor growth, but only one mouse treated with anti-4-1BB alone and none treated with anti-CTLA4 survived (Figure 4A, B). In contrast, approximately 60% of mice treated with ADWA-11 in combination with either anti-CTLA4 or 4-1BB were long-term survivors. As we found for the CCK168 and CT26 complete responders, re-challenge of all long-term survivors in the combined therapy groups showed minimal tumor growth followed by rapid tumor elimination, suggestive of long-term anti-tumor immunity (Figure 4C).

**Figure 4.**
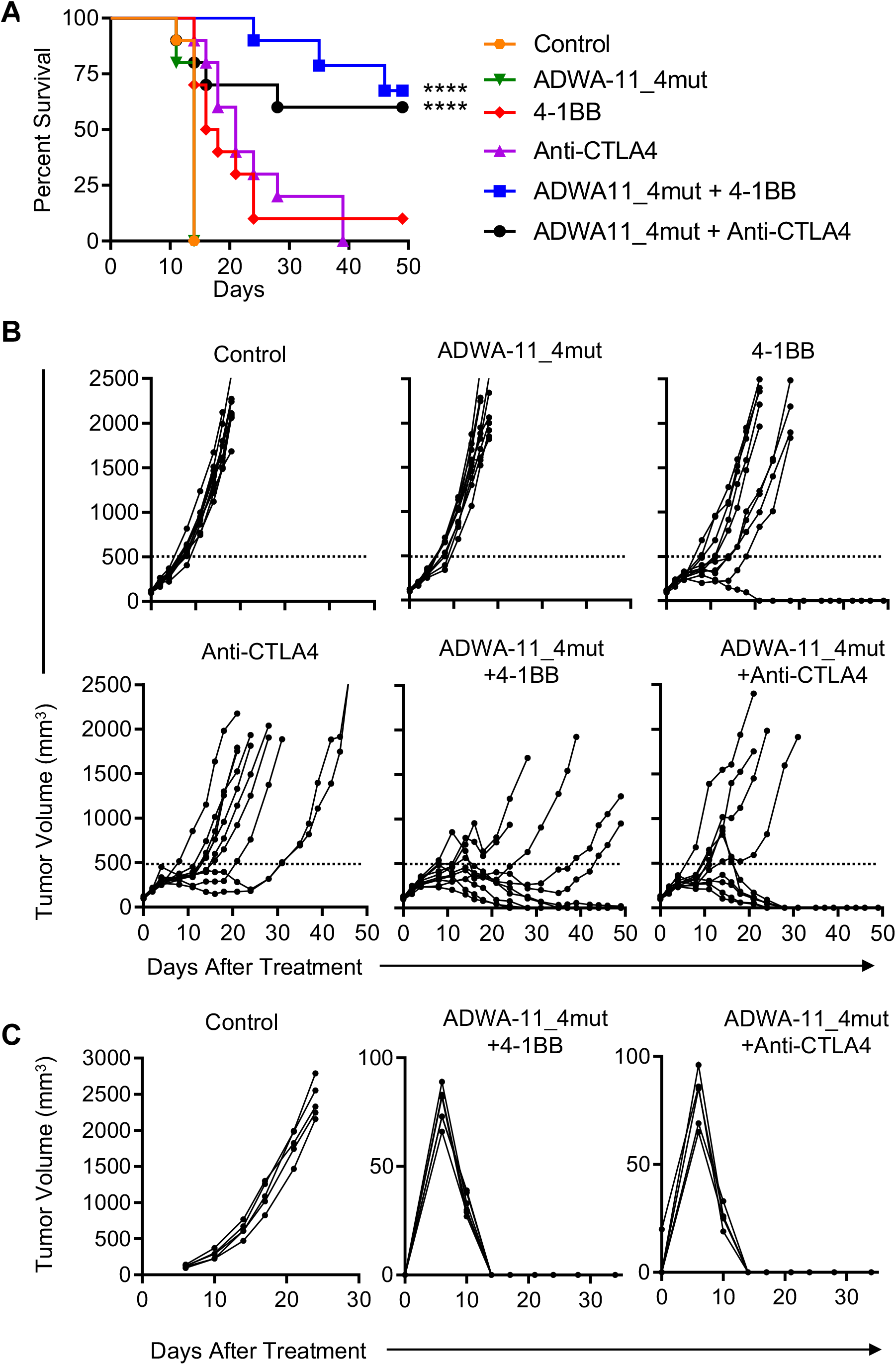
Combination treatment of ADWA-11_4mut with either the checkpoint inhibitor, anti-CTLA4 or the immune activator, 4-1BB, also improved survival, inhibited tumor growth and induced long-term anti-tumor immunity in the EMT6 mammary carcinoma model. (A) Survival curves and individual growth curves (B) in mice with orthotopic implantation of EMT6 cells treated with isotype control antibody, ADWA-11_4mut, 4-1BB, anti-CTLA4, combination of ADWA-11_4mut and 4-1BB, or combination of ADWA-11_4mut and anti-CTLA4. Data reported as percent survival, n=10 in each group. ****p<0.0001 by log-rank Mantel-Cox test. (C) Tumor re-challenge in mice that survived 50 days in the EMT6 model after treatment with the combination of ADWA-11_4mut and anti-CTLA4 or the combination of ADWA-11_4mut and 4-1BB. Control mice were not previously exposed to tumor. Re-challenged mice were assessed for 30 days. n= 5 control, n= 6 ADWA-11_4mut + anti-CTLA4, n=5 ADWA-11_4mut + 4-1BB

### ADWA-11 treatment inhibits TGFβ/Smad3 signaling in tumors and increases expression of genes normally repressed by TGFβ in CD8+ T cells

As noted above, all of the previously identified effects of tissue specific knockout of *Itgb8* could be explained by a loss of TGFβ signaling activity. To determine the contribution of αvβ8 to overall TGFβ activity in tumors, we measured the effects of ADWA-11 on phosphorylation of the early TGFβ signaling effector SMAD3 by western blotting of whole tumor lysates. ADWA-11 significantly inhibited pSMAD3 in CCK168, EMT6 and TRAMPC2 tumors, suggesting that αvβ8 is a major contributor to TGFβ activation in each of these tumor models (Figure 5A). Since our immunophenotyping and CD8+ depletion results suggested that CD8+ T cells are the functionally important target of ADWA-11 treatment, we sought to determine whether ADWA-11 might be acting by inhibiting TGFβ presentation to CD8+ T cells, with a consequent inhibition of TGFβ signaling in these cells. Previous work from Joan Massague’s lab(Thomas and Massague, 2005) identified a group of genes that were significantly repressed in vitro by activated CD8+ T cells treated with active TGFβ1. We used qPCR to interrogate expression of a panel of these genes (Granzyme B, Interferon-γ, Granzyme A, and Fas ligand) that we chose because each of them could play a role in enhanced CD8+ mediated tumor killing. Expression of all but one of these genes was significantly upregulated in CD8+ T cells sorted from CCK168 and EMT6 tumors (Fig. B,C), and all four genes were significantly upregulated in CD8+ T cells from ADWA-11 treated TRAMPC2 tumors (Figure 5D), suggesting that ADWA-11-mediated tumor suppression might, at least in part, be a consequence of inhibition of TGFβ presentation to CD8+ T cells.

**Figure 5.**
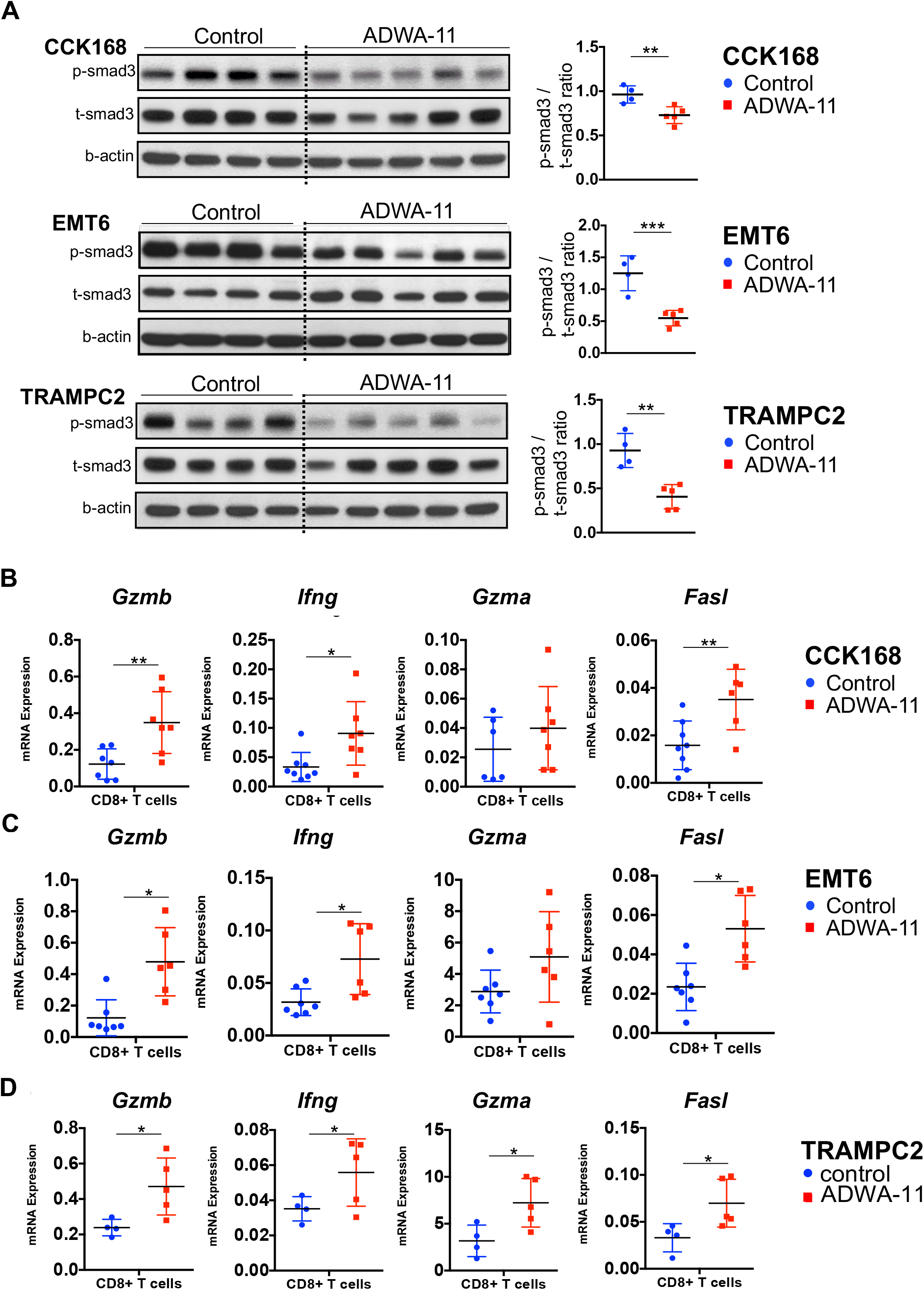
ADWA-11 inhibits TGFβ signaling in tumors and increases expression of genes known to be inhibited by TGFβ in CD8+ T cells. (A) Western blotting for pSMAD3 (and total SMAD3 and β-actin as loading controls) in whole tumor lysates from CCK168, EMT6 and TRAMPC2 tumors. mRNA expression of a set of genes known to be inhibited by TGFβ (Granzyme B, Interferon-γ, Granzyme A and Fas ligand) from intratumoral CD8+ T cells from control or ADWA11 treated mice harboring CCK168 (B) EMT6 (C) and TRAMPC2 (D) tumors. Data in graphs are mean ± S.D, n=5-8 per group. *p<0.05, **p<0.01 by t-test.

### Characterization of integrin αvβ8 expression on immune cells in the tumor microenvironment

Flow cytometry of in vitro cultures of each of the four tumor types studied showed a broad range of expression of αvβ8, with high expression in CCK168 and TRAMPC2 cells, minimal expression in EMT6, and undetectable expression in CT26 (Figure. S1D). Unlike the cases for cells in culture, none of the antibodies we evaluated against αvβ8 were able to reliably detect this integrin in cells disaggregated from in vivo tumors. We therefore used flow cytometry to sort a variety of cell types (Figure. S5) from three tumor types representing a range of αvβ8-expression on tumor cells (CCK168, EMT6 and TRAMPC2) and evaluated *Itgb8* mRNA expression by qPCR (Figure 6A). *Itgb8* expression was consistently seen at the highest levels on CD4+CD25+ T cells which, as expected, were the major sorted population expressing the Treg marker, *FoxP3* (Figure S5C). Even in CCK168 and TRAMPC2 tumors, where the tumor cells in vitro expressed easily detected surface expression of αvβ8, the level of *Itgb8* RNA expression was minimal in tumor cells harvested from in vivo tumors and more than 10 times higher in CD4+CD25+ T cells than in tumor cells. Importantly, *Itgb8* RNA expression was 3-5 fold higher in CD4+CD25+ T cells of the tumor compared to those isolated from spleen or tumor-draining lymph nodes (Figure S5C).

**Figure 6.**
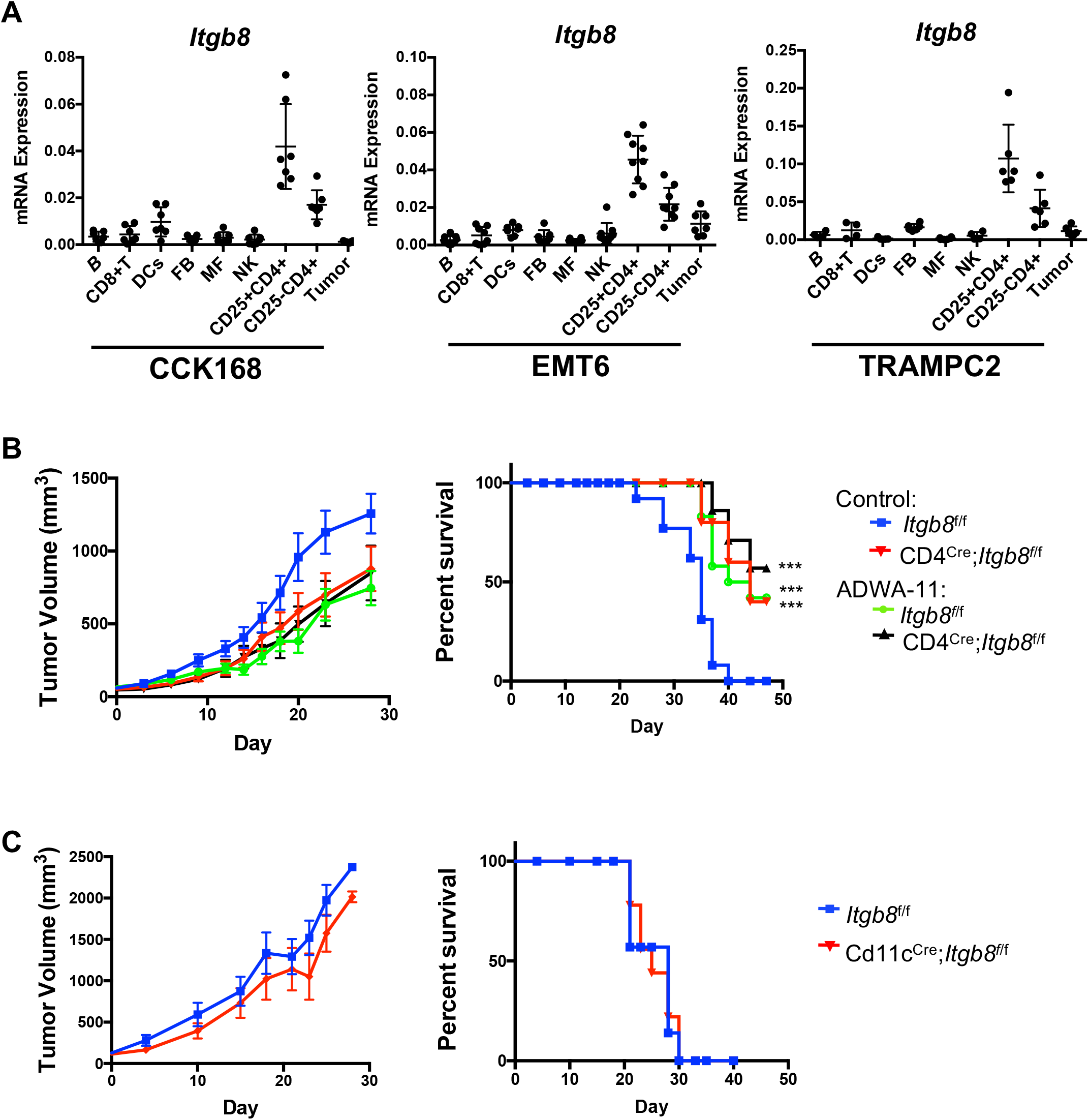
*Itgb8* is most highly expressed in CD25+CD4+ T cells and deletion of *Itgb8* specifically from T cells, but not from CD11c+ cells, inhibits in vivo growth of TRAMPC2 tumors. (A) Flow cytometry was performed to sort cell populations (Figure S5) from CCK168, EMT6 and TRAMPC2 tumors. mRNA was extracted and rt-PCR was performed to detect expression of integrin β8 by sorted cell populations. TRAMPC2 tumor cell lines were injected subcutaneously into CD4-Cre;*Itgb8*-f/f (B) or CD11c-Cre;*Itgb8-*f/f mice (C) or b8f/f littermate controls After tumors grew to 100-150mm^3^, mice were randomized and separated into two treatment groups for each genotype. Mice were injected i.p. with ADWA-11 or isotype control antibody on days 0 and 7. Tumor growth were plotted showing average volume of each group. Kaplan-Meier survival curves and individual growth curves of tumors were recorded and measured every other day. Survival curves used a tumor volume of 2,000 mm^3^ as cutoff. Mice were euthanized when tumors reached ≥ 2000 mm^3^, if extensive tumor ulceration was observed or at the 45-day endpoint. Data reported as percent survival, n=10-18 in each group. ***p<0.001 by log-rank Mantel-Cox test.

### Specific deletion of Itgb8 from T cells delays in vivo tumor outgrowth and is redundant to effects of ADWA-11 on tumor growth inhibition

Since we found high expression of *Itgb8* in CD4+CD25+ T cells, we next sought to determine if genetic deletion of *Itgb8* in T cells could recapitulate the suppressive effects of ADWA-11 on tumor growth. We generated CD4-Cre;*Itgb8-f/f* mice in which *Itgb8* is deleted in all T cells. TRAMPC2 tumor cells were injected into CD4-Cre;*Itgb8*-f/f and *Itgb8*-f/f littermates, and mice in both groups were untreated or treated with ADWA-11. Tumor growth was significantly suppressed (Figure 6B, S6A) and survival was enhanced in CD4-Cre;*Itgb8*-f/f mice compared to *Itgb8*-f/f littermates. As we previously found for control mice, ADWA-11 significantly inhibited growth and enhanced survival in Cre-negative *Itgb8*-f/f mice. However, ADWA-11 showed no additional benefit in CD4-Cre;*Itgb8*-f/f mice, strongly suggesting that αvβ8-mediated suppression of tumor immunity in this model is entirely due to αvβ8 expression by T cells. Because we and others have previously described functionally important effects of deletion of *Itgb8* from dendritic cells, we performed similar experiments in mice in which *Itgb8* was deleted from dendritic cells (CD11c-Cre;*Itgb8*-f/f) but found no effect of this deletion on tumor growth or survival (Figure 6C, S6B).

## Discussion

In this study, we show that ADWA-11, a blocking monoclonal antibody specific for the αvβ8 integrin, can inhibit tumor growth and enhance CD8+ T cell expression of a suite of genes involved in tumor killing in multiple syngeneic tumor models. ADWA-11 also dramatically enhances responses to multiple immunomodulators (anti-PD-1, anti-CTLA-4 blocking antibodies, and anti-4-1BB agonist antibodies) and to radiotherapy. Depletion of CD8+ T cells abrogates the beneficial effects of ADWA-11 and anti-PD-1 treatment suggesting that enhanced tumor cell killing by CD8+ T cells is critical for the anti-tumor effects of ADWA-11. Although in two of the tumor models we studied αvβ8 was expressed on the tumor cells prior to in vivo implantation, the other two responsive models expressed little or no αvβ8. In every model we examined, the highest level of *Itgb8* mRNA was found in CD4+CD25+ Treg cells. Furthermore, knockout of *Itgb8* specifically in T cells mimicked the effects of ADWA-11, suggesting that αvβ8 expressed on T cells is responsible for suppression of anti-tumor immunity. Since the major T cell subset we found to consistently express high levels of *Itgb8* in tumors was CD25+CD4+ cells, we suspect that these Tregs cells are responsible for this effect, although low levels of *Itgb8* expression in CD8+ and CD4+CD25-T cells may also contribute to TGFβ activation and consequent immunosuppression.

All previously described effects of loss of αvβ8 *in vivo* can be explained by contextual loss of activation of TGFβ, so we suspect that inhibition of TGFβ activation is responsible for these effects(Melton et al., 2010; Mu et al., 2002; Travis et al., 2007; Worthington et al., 2015). This hypothesis is supported by our observation that ADWA-11 significantly inhibited overall tumor TGFβ signaling in all of the models we studied. Our finding that a suite of genes involved in antigen-triggered cytotoxicity, including granzyme A and B, IFNγ and Fasl, which have all been previously shown to be inhibited by TGFβ in CD8+ T cells(Thomas and Massague, 2005), were coordinately increased in CD8+ T cells purified from tumors in mice treated with ADWA-11, suggests that TGFβ activated by αvβ8 expressed on CD4+ T cells inhibits tumor immunity at least in part by suppressing the cytotoxic program in CD8+ T cells. Short-term therapy with ADWA-11 led to long-term tumor suppression and resistance to subsequent re-challenge, strongly supporting the induction of long-term anti-tumor immunity.

One previous study described inhibition of tumor growth by treatment with an antibody against αvβ8, but the authors of that paper suggested that αvβ8 expression on tumor cells was critical for this effect(Takasaka et al., 2018). Most of the studies in that paper were based on a somewhat artificial system in which Lewis Lung Cancer cells or B16 melanoma cells were transfected to ectopically express αvβ8. The antibody used in that study had a wild type Fc domain, so could have been inhibiting tumor growth of transfected cells by ADCC. Because they did not detect αvβ8 expression by T cells using flow cytometry, they concluded that the integrin was only expressed on tumor cells. However, in this study, and in previously published studies showing functional effects of αvβ8 on Tregs(Stockis et al., 2017; Worthington et al., 2015), none of the available αvβ8 antibodies could detect αvβ8 by flow cytometry on Tregs. The basis for this limitation of antibodies for flow cytometry is not known, but we speculate that the epitopes recognized by these antibodies are not accessible on Tregs, perhaps because they are masked by other proteins (for example GARP) present in complex with αvβ8. Moreover, a recent study showed that integrin β8 can activate TGFβ without release of the mature ligand from the latent complex, that would also mask detection (Campbell et al., 2020). In the current study, we employed a range of tumor models, including some in which tumor cells expressed αvβ8 and others in which there was little or no detectible expression of either *Itgb8* mRNA or αvβ8 protein, and clearly show that αvβ8 blockade can be effective with or without αvβ8 expression on tumor cells. Our observations that deletion of *Itgb8* from T cells is as effective as systemic inhibition by ADWA-11, and our observation that administration of ADWA-11 to mice lacking *Itgb8* on T cells does not further inhibit tumor growth, strongly suggest that the effectiveness of αvβ8 blocking antibody is not simply due to blocking αvβ8 on tumor cells. Based on our results, we cannot be certain that αvβ8 on tumor cells does not contribute, in some circumstances, to suppression of anti-tumor immunity, but it is clear that much of the beneficial effects of αvβ8 blockade are due to inhibition of αvβ8 on T cells.

A previous study demonstrated combined effects of anti-PDL-1 and systemic blockade of TGFβ in the EMT6 model, and like us, also showed that CD8 depletion abrogated these effects and that the effect on tumor growth was associated with accumulation of Granzyme B-expressing, CD8+ T cells within the tumors. However, in that study, inhibition of TGFβ alone had no effect on either tumor regression or accumulation of Granzyme B-expressing CD8+ cells, whereas in the current study we saw clear effects of monotherapy with our αvβ8-blocking antibody. We think these differences may be due to the relative potency of the antibodies used in these different experiments. 1D11, the TGFβ inhibitory antibody used in the previous study, is not very potent, with an IC 50 ~ 40-fold lower than that of ADWA-11 for inhibition of TGFβ-activated by αvβ8 in a co-culture bioassay (Figure S1A). In our own experience, doses of this antibody as high as 30 mg/kg did not completely abrogate TGFβ signaling in vivo(Tsujino et al., 2017). In contrast, ADWA-11 is a highly potent antibody, with an IC50 for inhibition of TGFβ activation in the low nanomolar range (Figure S1). Furthermore, αvβ8 is not widely expressed, and when it is expressed is present at only several thousand receptors/cell. In contrast, TGFβ isoforms, all recognized by 1D11, are diffusely stored in the extracellular space in most tissues, making complete inhibition of these ligands difficult to achieve, and even more inaccessible if, as recently shown, integrins can activate TGFβ and its receptors without release of mature TGFβ from the latent TGFβ complex (Campbell et al., 2020). Of course, we cannot absolutely exclude the possibility that αvβ8 has effects on anti-tumor immunity that are independent of inhibition of TGFβ activation, but thus far no such effects have been convincingly demonstrated. In any case, our finding that inhibition of αvβ8 is at least as potent in enhancing anti-tumor immunity as previously reported global inhibitors of TGFβ make αvβ8 inhibitors attractive candidates to abrogate the potential toxicity of global inhibition of TGFβ.

In this report, we show that the αvβ8 integrin is expressed on CD25+CD4+ T cells in multiple syngeneic tumor models and is a potent modulator of the anti-tumor immune response. While efficacy was seen for monotherapy targeting this integrin in most of the tumor models we studied, efficacy can be substantially improved by combining inhibition of αvβ8 with either the checkpoint inhibitors, anti-PD-1 or anti-CTLA4, an agonist of the immune activator, 4-1BB, or by combining αvβ8 monotherapy with radiotherapy. Intriguingly, we found that *Itgb8* expression was much higher on intratumoral CD4+CD25+ T cells than those isolated from draining lymph nodes and spleen, a feature that might benefit immune-mediated tumor rejection by this drug whilst minimizing harm to normal tissues that might arise from reduced peripheral tolerance. Taken together, these results identify the αvβ8 as a promising target for tumor immunotherapy, even in tumors that do not express this integrin on tumor cells.

## Acknowledgments

**Funding:** This work was supported by a grant from the joint UCSF-Pfizer Centers for Translational Innovation (CTI) to DS and RJA, NCI R01CA197198 (RJA) and a NCI CTD^2 grant, U01CA217864 (PI, William Weiss). The work was supported by the use of UCSF HDFCCC shared resource facilities, Laboratory of Cell Analysis and Preclinical Therapeutics Core. JTL was supported by NIH KHL111208A, DSM was supported by a fellowship from the Swiss National Science Foundation. **Authors’ contributions:** H-YM and EDM wrote the manuscript, designed, performed experiments. BL and JBL designed and performed experiments and oversaw immunophenotyping. DSM, BCH performed *in vivo* tumor experiments. KHS, XR, TA, NIR, JSDC, TT performed ADWA-11 antibody characterization experiments and designed, performed and interpreted immunostaining experiments and in vivo tumor experiments. EDM, LP, BH and MA performed in vivo tumor experiments with CCK168. BZ, MDR, MBH and EDM performed mouse immunophenotyping experiments. SY, AG, KN, BY, KN, JDP developed effectorless ADWA-11 antibody, performed *in vivo* tumor experiments using effectorless ADWA-11 antibody and analyzed data. AA developed ADWA-11 antibody, designed and performed experiments. RJA, DS wrote and edited manuscript, designed experiments, analyzed data and oversaw all aspects of the work described. **Competing interests:** DS and RA were funded by a grant from the joint UCSF-Pfizer CTI. Sharon Yang, Anand Giddasbasappa, Kavon Koorbehesht, Bing Yang, Kyle Niessen and Joseph Dal Porto are employees of Pfizer. UCSF and Pfizer have filed a joint patent covering the use of a humanized version of ADWA-11 for cancer immunotherapy.

## Methods

### Animals

Mice were bred and maintained according to approved protocols by the University of California, San Francisco, or Pfizer Inc., Institutional Animal Care and Use Committee. Wild type FVB mice were purchased from Jackson Laboratories (The Jackson Laboratories, stock #001800). Wild type Balb/c mice were purchased from Charles River Laboratories (Charles River Laboratories, strain code 028). Wild type C57/B6 mice were purchased from Jackson Laboratories (The Jackson Laboratories, stock #001800).

### Tumor models

CCK168 cells, a chemically induced squamous cell carcinoma cell line derived from FVB mice^6^, were injected subcutaneously in the right flank region in syngeneic wild type FVB mice (1.5×10^4^ cells/mouse). Tumors were allowed to grow over 14 days. Mice selected for the experiment had tumor size at least 0.5 cm in diameter and were randomized to different antibody treatment groups using a random number generator (https://www.random.org). The study team was blinded to the treatment groups of 10 mice per group. Mice were weighed daily, and tumor size was measured every other day for the duration of the study using a traceable digital caliper (Fisher Scientific, model #14-648-17). Mice were euthanized when tumor size exceeded 2000 mm^3^ or developed a large ulceration at the tumor site. Appropriate antibody for each group and isotype control antibodies were injected on days 0, 4, and 8 of the experiment for anti-PD-1 and day 0 and 7 for ADWA-11. Doses of intraperitoneal antibodies injected were ADWA-11 (10 mg/kg), Anti-PD-1 (10 mg/kg), control antibody ADWA-21 (for ADWA-11, 10 mg/kg), and control 2A3 (for PD-1, 10 mg/kg). Control ADWA-21 binds only human integrin-β8, but not mouse integrin-β8. For CD-8 depletion studies, intraperitoneal anti-CD-8 (Bio X Cell® BE0004-1 Clone 53-6.72) or control antibody was injected at a dose of 10 mg/kg on days 0, 4, and 8. Combined ADWA-11 (10 mg/kg) and anti-PD-1 (10 mg/kg) were injected on days 1, 5, and 9 in the CD8 depletion studies.

EMT6 (ATCC®, CRL2755™) cells, a mouse epithelial mammary carcinoma cell line, were injected into the fourth mammary fat pad (5×10^4^ cells/mouse) of Balb/c mice, and mice were followed daily until the tumors reached 0.5 cm for inclusion in the study(Zhang et al., 2016). Subsequent treatment regimens and measurements were performed as above.

TRAMPC2 (ATCC®, CRL2731™) cells, a mouse epithelial prostate adenocarcinoma cells were injected subcutaneously in the right flank region in syngeneic wild type C57/B6 mice (1×10^6^ cells/mouse with matrigel). Tumors take about four weeks to reach 100mm^#^ for the treatment that described as above.

CT26 mouse colon carcinoma cells (ATCC® CRL-2638™, 2.5 × 10^5^ cells/mouse) were injected into the subcutaneous flank of female Balb/c mice (Charles River Labs). Tumors were allowed to grow to 50-100 mm^3^ in size for inclusion in the study. For these studies, ADWA-11_4mut or isotype control 2B8_mIgG_4mut were injected on days 0, 4, 8, 12, anti-PD-1 (RMP1-14, BioXcell) or isotype control 2A3_rat IgG (BioXcell) were injected on days 0, 4, 8 through an intravenous route. All antibodies were dosed at 10mg/kg. All mice on Day 5, except the no treatment control group, were Exposed to tumor targeted 5 Gy dose of radiation. Tumor growth was measured twice per week with digital calipers and reported as volume (length × width × width × 0.5). For the re-challenge experiment, on day 51 (post first antibody treatment) mice with a complete response and naïve mice were implanted on the contralateral flank with 2.5 × 10^5^ CT26 cells in PBS and tumor growth was monitored as described above.

ADWA-11_4mut was also tested in the EMT-6 tumor model. 1 × 10^6^ EMT6 cells (ATCC®, CRL2755™) were injected into the fourth mammary fad pad of female Balb/c mice (Charles River Labs). Tumors were allowed to grow up to 50-100 mm^3^ in size. Mice were randomized into antibody treatment groups and operators were blinded to treatment groups. ADWA-11_4mut 10mg/kg or control 2B8_mIgG4mut 10mg/kg, anti-CTLA4 (9D9 BioXcell) or isotype control E.tenella-mIgG2b 10mg/kg were injected on days 0, 4 and 8, and 4-1BB (MAB9371, R&D systems) 1mg/kg was injected on days 0 and 4 through an intravenous route. Tumor growth was measured twice per week with digital calipers and reported as volume (length × width × width × 0.5). For the re-challenge experiment, on day 51 (post first antibody treatment) mice with a complete response and naïve mice were implanted in the contralateral fat pad with 1 × 10^6^ EMT-6 cells and tumor growth was monitored as described above.

### Generation of avβ8 blocking antibody ADWA-11

Integrin β8 knockout mice, crossed into the outbred CD1 background to permit post-natal survival were immunized with recombinant αvβ8 integrin (R&D Systems, 4135-AV-050) 50μg per mouse every two weeks. Sera from immunized mice were screened by solid phase immunoassays and used to identify mice for hybridoma generation. Antibodies from the hybridomas were labeled with APC fluorophore and further characterized by flow cytometry using SW480 cells, which normally do not express any αv integrins except αvβ5, were transfected to express integrin αvβ8 or αvβ3 or αvβ6 as negative controls. We performed flow cytometry on each cell line using labelled ADWA-11 or antibodies to αvβ5 (Alula) or αvβ3 (Axum-2) or αvβ6 (10D5)(Huang et al., 1998; Su et al., 2012; Su et al., 2007). Cell adhesion assays were performed with U251 cells that express integrin αvβ8 on dishes coated with TGFβ1 latency associated peptide 1μg/ml(Kueng et al., 1989). Blockade of TGFβ activity was determined by TMLC luciferase assay (mink lung epithelial cells expressing a TGFβ sensitive portion of PAI-1 promoter driving firefly luciferase expression)(Abe et al., 1994).

### Generation of effectorless ADWA-11 antibody (ADWA-11_4mut)

To construct an anti-αvβ8 antibody with reduced Fc receptor binding and effector function the variable regions from ADWA11 were subcloned into a mIgG1 Fc backbone that contained E233P, E318A, K320A, and R322A mutations, as outlined in https://www.google.com/patents/US20090155256. ADWA-11_4mut binding to mouse αvβ8 was assessed using C8-S mouse astrocyte cells (ATCC) and blockade of TGFβ activation was determined using C8-S cells in a co-culture TMLC luciferase assay, described above.

### Flow cytometry

Subcutaneous tumors were isolated from the mice using scissors and blunt dissection. The tumors were placed in a petri dish with digestion cocktail of Collagenase XI (Sigma C9407) 2 mg/mL, Hyaluronidase (Sigma H3506) 0.5 mg/ML, and DNase (Sigma DN25) 0.1 mg/mL prepared in C10 media (RPMI 1640, Hepes 1%, Penicillin/Streptomycin 1X, fetal calf serum 10%, sodium pyruvate 1 mM, non-essential amino acids 1X, and beta-mercaptoethanol 0.45%). Tumors were minced using sterile scissors. The resultant slurry of cells was transferred into 50 mL conical tubes (Fisher Scientific #14-432-22) and the petri dish used to mince the tumor was rinsed with 2 mL C10 media to capture remaining cells. Cells were incubated in a shaker at 255 rpm for 45 minutes at 37°C. After incubation, 15 mL of C10 media was added to the digested tumor cells and gently vortexed for 15 seconds. The cell slurry was passed through a 100 μm mesh strainer (Falcon® #352360) into a clean 50 mL conical tube. Cells were pelleted by centrifugation for 5 minutes at 200 g at 4°C and reconstituted in PBS. Cell counts were performed using a hemocytometer (Fisher Scientific #02-671-6). Isolated single cell preps were used for cell surface and intracellular staining. After counting, 10 × 10^6^ cells were transferred into each well of a v-shaped 96-well plate for staining. Live dead staining with Ghost Dye™ Violet 510 (TONBO bioscience#13-0870) at 1:1000 for 20 minutes at 4°C. Fc receptor and non-specific binding was blocked with anti-CD16/30 (eBioscience #14061) for 10 minutes at 4°C. Surface staining was performed for 20 minutes at 4°C. For intracellular staining, cells were incubated in Fix/Perm buffer (eBioscience #88-8824) for 20 minutes at room temperature followed by intracellular cytokine staining with antibody cocktails for 20 minutes at 4°C. After completion of staining, cells were transferred into flow cytometry buffer (PBS with 2%FBS, Penicillin/Streptomycin/Glutamate, EDTA 2 mM) for analysis.

Antibodies used for intracellular cytokine staining experiments: CD45 AF700 (Invitrogen#56-0451-80), CD3e BV711 (Biolegend#100241), CD25 BV605 (BioLegend#102036), CD4 APC-Cy7(Invitrogen#47-0042-82), CD8a PE Cy7 (eBioscience#25-0081-81), NKG2A/C/E APC (Biolegend 564383), Granzyme-B PE (eBioscience#12-8898-82), FoxP3 PB-e450 (eBioscience#48-5773-82), IFNg BV650 (BioLegend#505831)

For cell stimulation, cells were stimulated prior to cell surface and intracellular staining. Approximately 3×10^6^ cells in 200μl of C10 media per well were incubated in round-bottom 96-well plates overnight in a tissue culture incubator in 5% CO2 at 37°C. Stimulation cocktail (Inomycin, PMA, Brefeldin-A, and Monensin 500x stimulation cocktail Tonbo #TNB4975-UL100) was added to cells which were incubated in tissue culture incubator in 5% CO2 at 37°C. for 4 hours. Cells were transferred to v-bottom wells for staining as outlined above.

Antibodies used for cell sorting experiments (Lymphocyte panel) : CD45 AF700 (Invitrogen 56-0451-80), B220 PerCP Cy5.5 (eBiosciences 45-0452-80), CD8 PE(eBiosciences 12-0081-81), CD4 APC-Cy7(Invitrogen 47-0042-82), NKG2A/C/E APC (Biolegend 564383), CD3e BV711 (Biolegend 100241), CD25 BV605 (BioLegend#102036)

Antibodies used for cell sorting experiments (Myeloid panel): CD45 AF700 (Invitrogen 56-0451-80), CD64 FITC (BioLgend#139307), Cd11b PE-Cy7 (BioLegend#101215), Thy1.2 BV421 (BioLegend#105341), Thy1.1 BV421 (BioLegend#202529), (Cd11c BV605 (BioLegend#117334), MHCII PE/Dizzle(BioLegend#107648). Flow cytometry was performed using a Attune NxT Flow Cytometer ™ (ThermoFisher). Cell sorting was sperformed using a BD FACSAria3™ (BD Biosciences) and analyzed using FlowJo™ (Tree Star Inc.).

### Quantitative PCR Analysis

Total RNA was isolated from sorted tumoral cells using Trizol Reagent (Life Technologies) according to the manufacture’s instruction. RNA was reverse transcribed by SuperScript IN VILO master mix(Invitrogen) with DNase treatment. Power SYBR Green master mix(Life Technologies) was used for quantification of cDNA on QuantStudio 5 Real Time PCR system (Thermo-Fisher). Primers used were listed in Table 1.

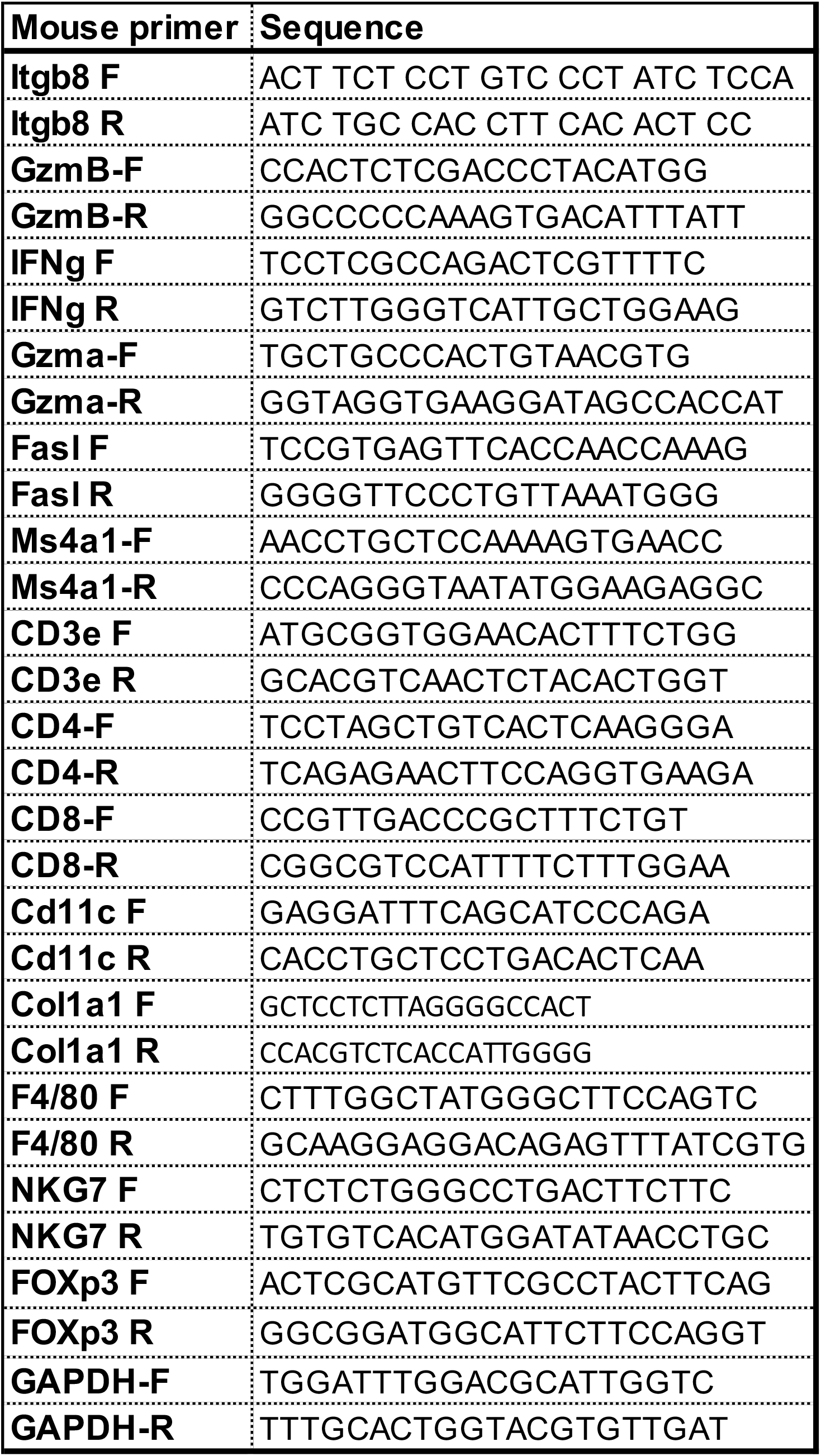

### Western Blotting

Tumors were harvested and homogenized in 500ul RIPA lysis buffer (50 mM Tris-HCl, pH 8, 150 mM NaCl, 1% NP-40, 0.1% SDS, 0.5% sodium deoxycholate) containing protease and phosphatase inhibitors (ThermoFisher Scientific). Tumor lysate protein concentration was measured using Pierce BCA Protein Assay (ThermoFisher Scientific). Tumor lysate then boiled at 95 C on a heat block in Laemmli sample buffer (4% SDS, 10% 2-mercaptoethanol, 20% glycerol, 0.004% bromophenol blue, 0.125 M SDS, pH 6.8), separated by electrophoresis on a 10% polyacrylamide gel, and transferred to a polyvinylidene difluoride membrane (Millipore Sigma). The primary antibodies and dilutions were as follows: anti-rabbit pSMAD3 Ab (Abcam ab52903 1:1000), tSMAD3(Abcam ab40854 1:1000) and anti-β-actin Ab (Sigma, a5441, 1:5000). Secondary antibodies were horseradish peroxidase (HRP) conjugated goat anti rabbit (Cell Signaling Technology, 1:3000) or goat anti mouse (Cell Signaling Technology, 1:5000). Chemiluminescent HRP substrate (PerkinElmer) was used for membrane development. Immunoreactive bands were visualized with a Mini-Med 90 X-Ray Film Processor (AFP Imaging). a was performed using ImageJ software.

### Immunostaining

Tumors were harvested and fixed in 4% paraformaldehyde at 4°C for 3 hours. The fixed tumors were immersed in 30% sucrose solution at 4°C overnight and embedded in O.C.T. compound (Tissue Tek®#4583). Frozen sections were stained by previously described protocols(Henderson et al., 2013; Rock et al., 2011). Antibodies used for immunostaining: FITC-conjugated or Alexa Fluor 594-conjugated anti-mouse CD8 antibody (BioLegend, clone 53-6.7). Confocal microscopy was performed on a Zeiss LSM 780 microscope.

### Statistical Analysis

One-way ANOVA was applied to compare three or more groups and *post-hoc* Tukey-Kramer tests were used to identify specific differences. Unpaired two tail T-test was used to compare two groups to determine significant change. Values are reported as mean ± standard division (S.D.). In survival studies, comparison of survival curves were calculated based on Log-rank Mantel-Cox test. Statistical analysis was performed using GraphPad Prism 5.0, Graphpad Software, San Diego, California, USA.

