## Supplementary material for "Integrin αvβ8 on T cells is responsible for suppression of anti-tumor immunity in multiple syngeneic models and is a promising target for tumor immunotherapy": Suppl Legend

### Supplemental Legend

#### Supplemental Figure 1. Characterization of ADWA-11

(A) TMLC cell co-culture bioassays performed with concentrations of ADWA-11 ranging from 0.067 to 66.67 pM and 1D11 ranging from 0.667 to 666.7 pM. TGFβ activity is reported as relative luciferase units based on PAI-1 luciferase reporter activity. n=3 per ADWA-11 and 1D11 dose, repeated 3 times.

(B) Cell adhesion assays performed on dishes coated with the latency associated peptide (LAP) of TGFβ1 in the presence of ADWA-11 in concentrations from 0.067 to 66.67 pM Adherent cells were stained with crystal violet and adhesion expressed as absorbance at 595 nm. n=3 per ADWA-11 dose, repeated 3 times.

(C) Antibodies to integrin αvβ3, αvβ5, αvβ6, and αvβ8 were used for flow cytometry of wild type colon carcinoma cells, SW-480, that do not express any of these integrins or SW-480 cells transfected to express β3 (SW-itgb3), β6 (SW-itgb6), or β8 (SW-itgb8). Representative flow cytometry plots are shown for each antibody and cell type tested.

(D) Live single cells were analyzed for αvβ8 expression with ADWA-11 in the dish cultured different tumors (CCK168, EMT6, TRAMPC2 and CT26).

**Supplemental Figure 2 Gating strategy for intracellular cell flow cytometry.**

Single cells were first gated for forward and side scatter to enrich for lymphocytes, then gated for live cells. Live cells were then positively gated for CD45+CD3+ immune cells and further gated for CD4+ and CD8+ cells. Intracellular signal for FoxP3 and IFNg were analyzed in the CD4+ population, and for Gzmb and IFNg were analyzed in the CD8+ population.

**Supplemental Figure 3. CD8+ T cell depletion within CCK168 tumors abrogates the anti-tumor effects of combinatorial ADWA-11 and anti-PD-1 therapy.**

(A) Immuno-depletion of CD8+ T cells. Micrographs show immunostaining with anti-CD8 (Green) counterstained with DAPI (blue) in CCK168 tumors isolated following combinatorial anti-PD-1/ADWA11 therapy with or without prior treatment with anti-CD8 depleting antibody or isotype control antibody. (B) Average tumor growth curves for CCK168 tumors pretreated with anti-CD8 depleting antibody or isotype control antibody one-day prior to ADWA-11/anti-PD-1 combination therapy.

**Supplemental Figure 4. ADWA-11_4mut therapy synergizes with anti-PD-1 to improve survival.**

Survival curves for mice implanted with subcutaneous CCK168 cells treated with isotype control antibodies, anti-PD-1, ADWA-11_4mut, or a combination of anti-PD-1 and ADWA-11_4mut.

**Supplemental Figure 5. Gating strategy for cell sorting and identification of tumor infiltrating immune cells and monocytes.**

**(A)** Single cells were first gated for forward and side scatter to enrich for lymphocytes, then sorted with a Live CD45+ gate and then positively gated for CD3+NKG2A/C/E- (CD3 cells), CD3- NKG2A/C/E+ (NK). Then CD3 population were positively gated for CD8+CD4- (CD8+ T cells) and CD8-CD4+ (CD4+ cells). B220+ (B cells) were gated from CD3- NKG2A/C/E- cells. CD4+ cells were further gated for CD25+CD4+ and CD25-CD4+ cells.  Single cells were then gated for forward and side scatter to enrich for myeloid cells. After dumping out dead cells, live cells were gated for CD45-Thy1- (tumor cells), Thy1+CD45- (fibroblasts)(FB). CD45+Thy1- cells were then gate for CD64+CD11b+ (macrophages)(MF). CD64- cells were further gated for MHCII$^{hi}$Cd11c$^{hi}$ (dendritic cells)(DC). (B) Expression of population marker genes were tested by qPCR to determine the sorted cell population.  Itgb8 expression were found highly expressed in sorted CD4+ cells. (C) Expression of Foxp3 was detected by qPCR to determine enrichment of regulatory T cells in sorted CD25+CD4+ cells. CD25+CD4+ cells express higher Itgb8 in tumor than lymph node or spleen.

**Supplemental Figure 6. Deletion of itgb8 specifically from T cells, but not from CD11c+ cells, inhibits in vivo growth of TRAMPC2 tumors**

TRAMPC2 tumor cell lines were injected subcutaneously into CD4-Cre;*Itgb8*-f/f (A) or CD11c-Cre;*Itgb8*-f/f mice (B) or b8f/f littermate controls. Mice were injected i.p. with ADWA-11 or isotype control antibody on days 0 and 7. Individual growth curves of tumors measured every other day for each group. Mice were euthanized when tumors reached ≥ 2000 $mm^3$, if extensive tumor ulceration was observed or at the 45-day endpoint.
