## Supplementary figures and images for "Integrin αvβ8 on T cells is responsible for suppression of anti-tumor immunity in multiple syngeneic models and is a promising target for tumor immunotherapy"

### Suppl Fig 1

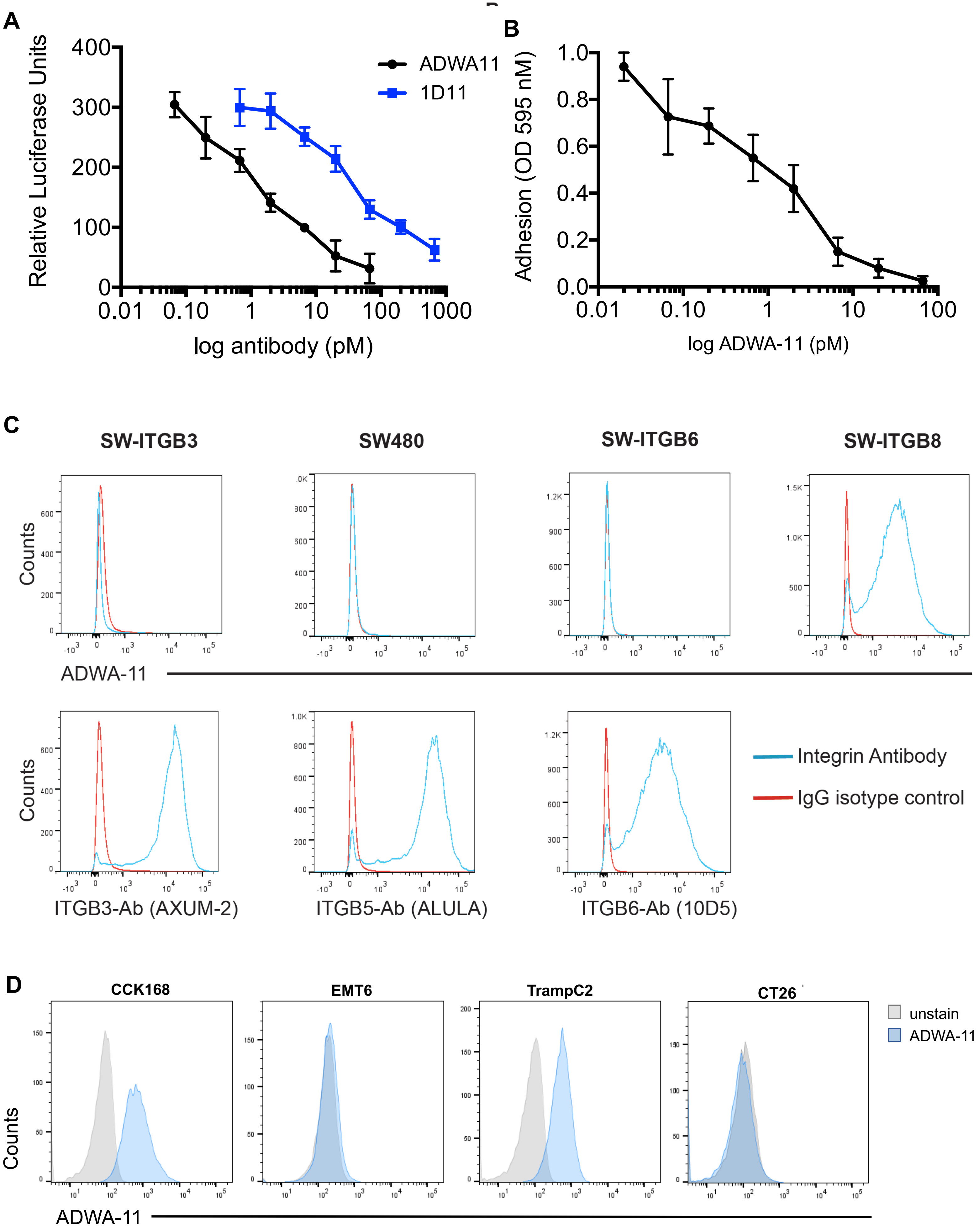

### Suppl Fig 2

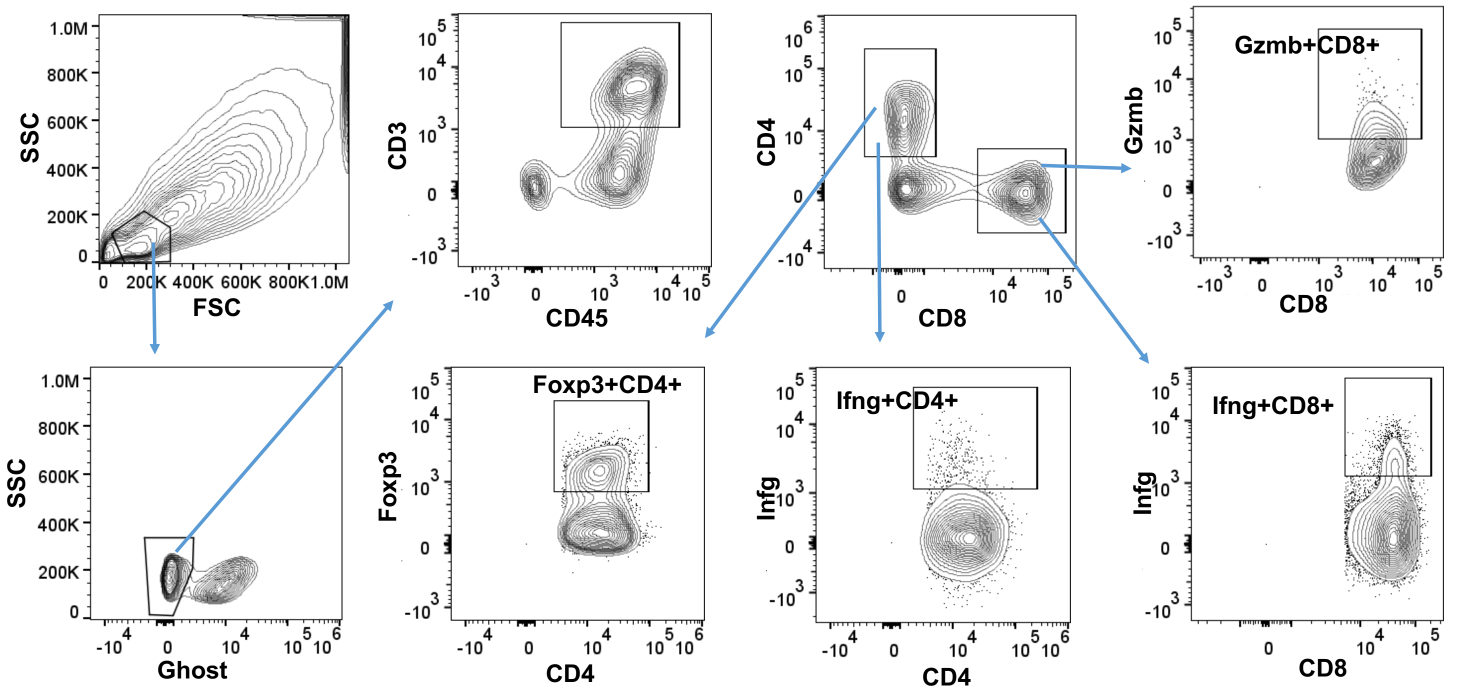

### Suppl Fig 3

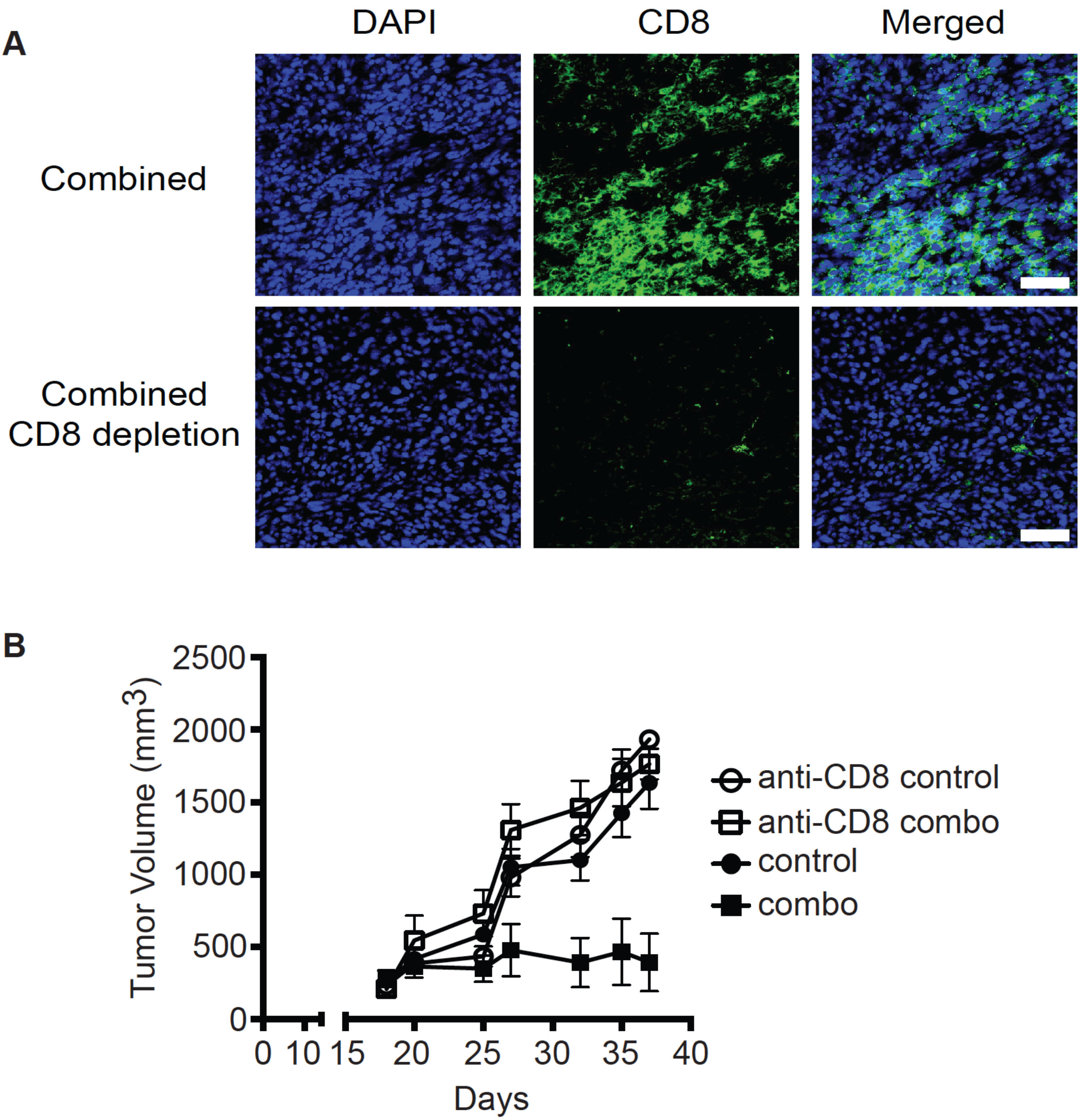

### Suppl Fig 4

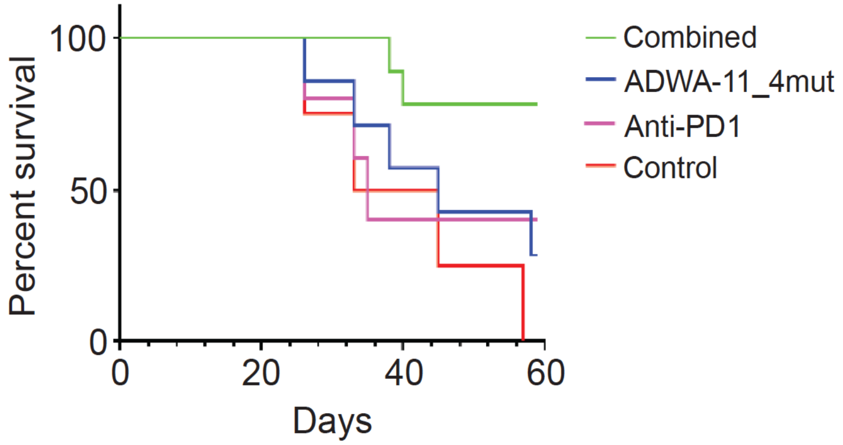

### Suppl Fig 5

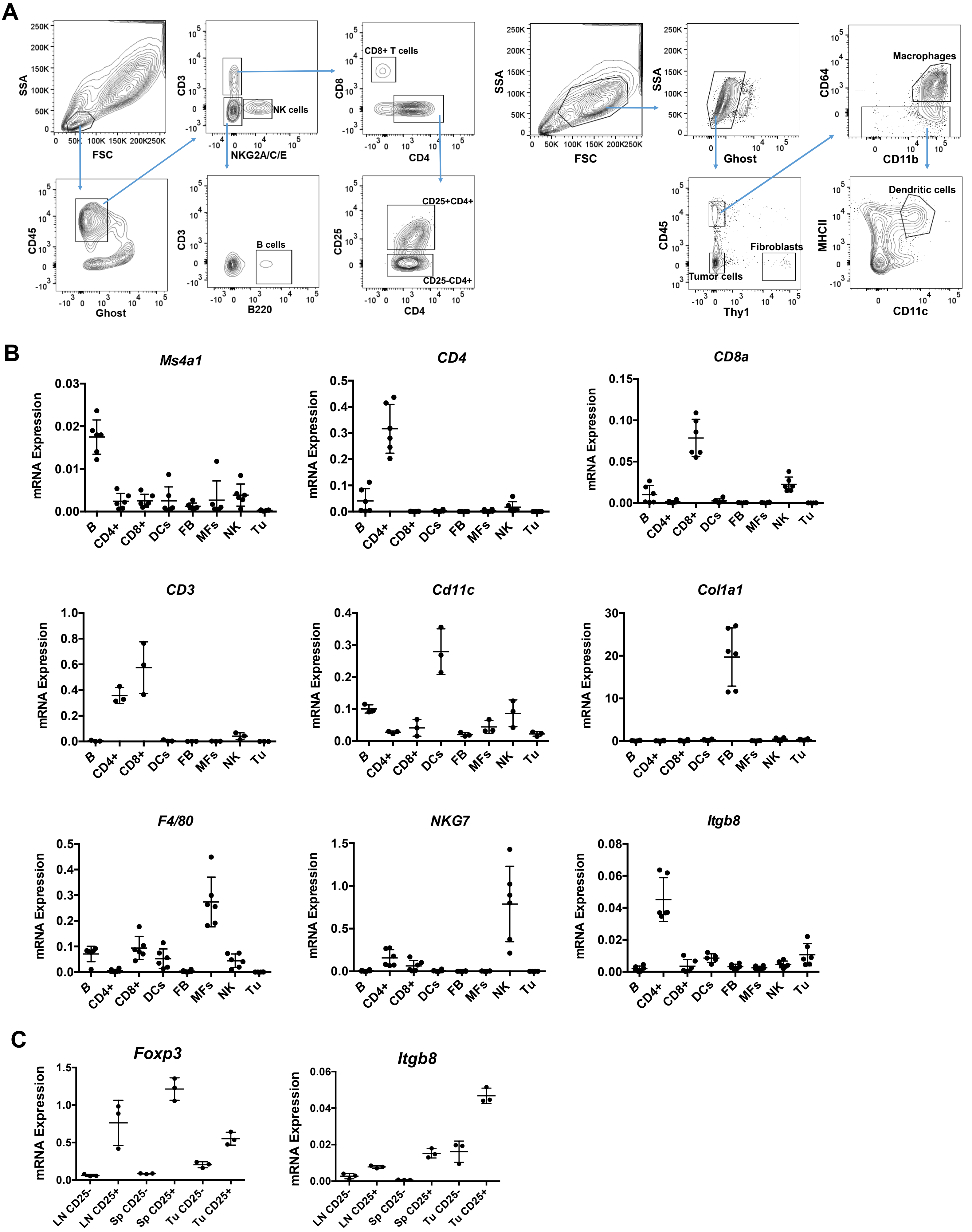

### Suppl Fig 6

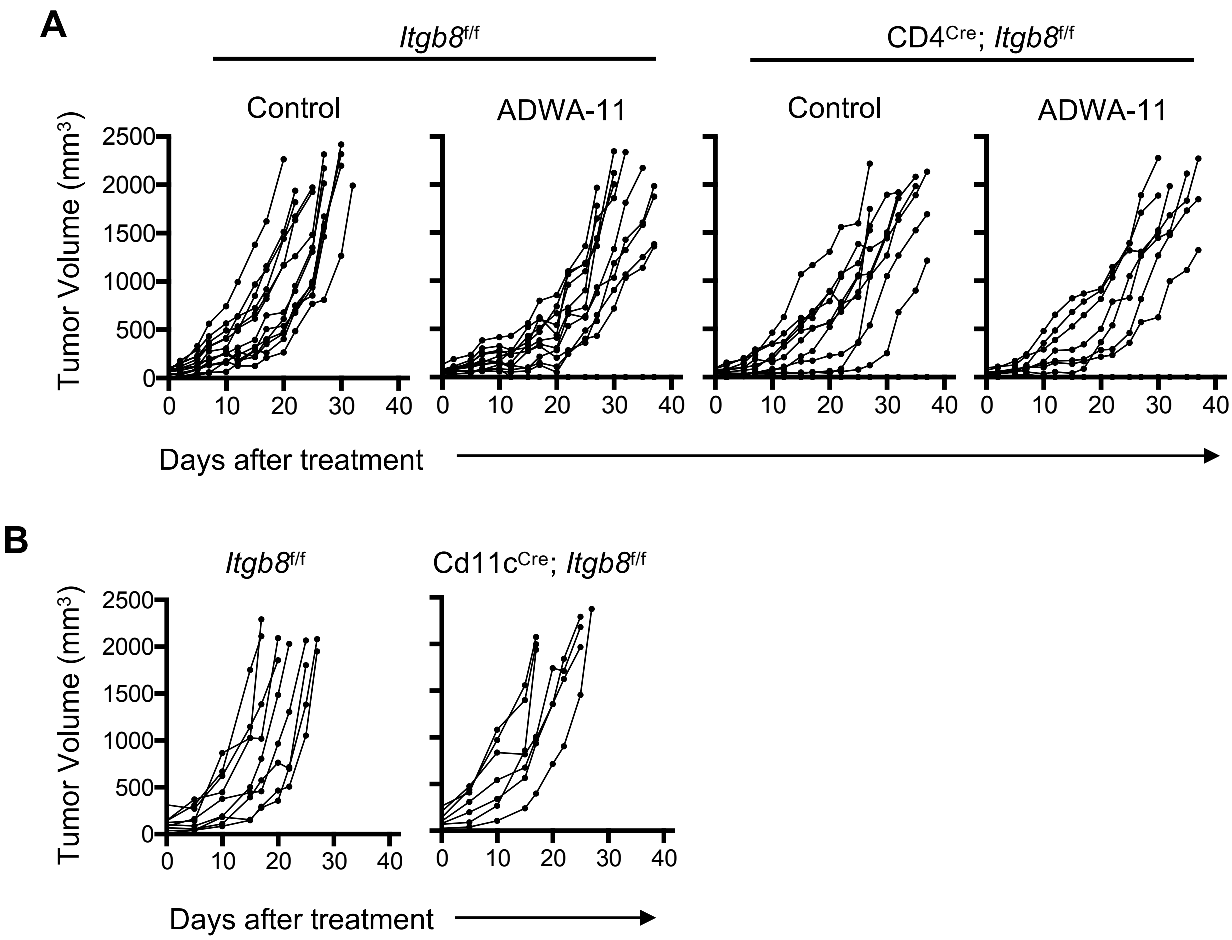
